# Long-Read RNA-sequencing reveals transcript-specific regulation in human-derived cortical neurons

**DOI:** 10.1101/2024.11.08.622610

**Authors:** Jishu Xu, Michaela Hörner, Elena Buena Atienza, Kalaivani Manibarathi, Maike Nagel, Stefan Hauser, Jakob Admard, Nicolas Casadei, Stephan Ossowski, Rebecca Schüle

**Affiliations:** Centre for Neurology and Hertie Institute for Clinical Brain Research, University of Tübingen, Tübingen, Germany; Institute of Medical Genetics and Applied Genomics, University of Tübingen, Tübingen, Germany; Graduate School of Cellular and Molecular Neuroscience, University of Tübingen, Tübingen, Germany; Division of Neurodegenerative Diseases and Movement Disorders, Department of Neurology, Heidelberg University Hospital and Faculty of Medicine, Heidelberg, Germany; German Centre for Neurodegenerative Diseases (DZNE), Tübingen, Germany; NGS Competence Center Tübingen (NCCT), University of Tübingen, Tübingen, Germany; Institute for Bioinformatics and Medical Informatics (IBMI), University of Tübingen, Tübingen, Germany; Interdisciplinary Center for Neurosciences (IZN), Heidelberg, Germany; Faculty of Biosciences, University of Heidelberg, Germany

**Author notes:** Corresponding author: Prof. Dr. med. Rebecca Schüle, Division of Neurodegenerative Diseases and Movement Disorders Department of Neurology, Heidelberg University Hospital and Faculty of Medicine Im Neuenheimer Feld 400, 69120 Heidelberg Germany. These authors contributed equally to this study (Jishu Xu, Michaela Hörner).

**Keywords:** Long-read RNA-sequencing, transcriptomics, transcript usage, alternative splicing, human-derived cortical neurons, induced pluripotent stem cells

## Abstract

Long-read RNA sequencing has transformed transcriptome analysis by enabling comprehensive mapping of full-length transcripts, providing an unprecedented resolution of transcript diversity, alternative splicing, and transcript-specific regulation. In this study, we employed nanopore long-read RNA sequencing to profile the transcriptomes of human fibroblasts, induced pluripotent stem cells, and stem cell-derived cortical neurons, identifying extensive transcript diversity with 15,072 transcripts in stem cell-derived cortical neurons, 13,048 in fibroblasts, and 12,759 in induced pluripotent stem cells. Our analyses uncovered 35,519 differential transcript expression events and 5,135 differential transcript usage events, underscoring the complexity of transcriptomic regulation across these cell types. Importantly, by integrating differential transcript expression and usage analyses, we gained deeper insights into transcript dynamics that are not captured by gene-level expression analysis alone. Notably, differential transcript usage analysis highlighted transcript-specific changes in disease-relevant genes such as *APP, KIF2A*, and *BSCL2*, associated with Alzheimer’s disease, neuronal migration disorders, and degenerative axonopathies, respectively. This added resolution emphasizes the significance of transcript-level variations that often remain hidden in traditional differential gene expression analyses.

Overall, our work provides a framework for understanding transcript diversity in both pluripotent and specialized cell types, which can be used to investigate transcriptomic changes in disease states. Additionally, this study underscores the utility of differential transcript usage analysis in advancing our understanding of neurodevelopmental and neurodegenerative diseases, paving the way for identifying transcript-specific therapeutic targets.

## Introduction

Cellular differentiation is a complex process where cells transition from a pluripotent state to specialized mature phenotypes. This developmental continuum, from pluripotent stem cells to differentiated cell types like fibroblasts or neurons, is critical for understanding developmental biology, disease mechanisms, and providing therapeutic innovations (Mallon et al., 2013; Tanabe et al., 2015; Vierbuchen & Wernig, 2011). Robust cell systems have been developed to generate various cell types *in vitro*, including different types of neurons (Fowler et al., 2020; Gopalakrishnan et al., 2017; Ng et al., 2021; Pawlowski et al., 2017). Transcriptome studies in most neuronal models, however, have been limited to gene expression studies, which often provide only broad regulatory patterns. These analyses tend to overlook the complexity introduced by alternative splicing (AS)—a process that affects nearly 95% of human pre-mRNAs (Pan et al., 2008). AS facilitates differential gene expression across tissues and developmental stages, and the generation of diverse transcript variants, enabling differential gene expression across tissues and developmental stages (Baralle & Giudice, 2017).

The regulation of AS involves a complex network in which RNA-binding proteins (RBPs) act as cis-regulators by directly interacting with intronic and exonic splice enhancer and silencer sequences. AS is also controlled by trans-regulators, such as transcription factors (TFs), which modulate the transcriptional elongation rate and modify chromatin structure (Kornblihtt et al., 2004; Mohan et al., 2024; Nazim et al., 2024; Ng et al., 2021; Porter et al., 2018; Schmok et al., 2024; Schwartz et al., 2009). AS is particularly prevalent in the vertebrate brain, where it plays an essential role in key processes such as neurogenesis, synaptogenesis, axon guidance, and neural plasticity (Furlanis & Scheiffele, 2018; Nazim et al., 2024; Papadimitriou & Thomaidou, 2024; Su et al., 2018; Wang et al., 2024; Weyn-Vanhentenryck et al., 2018; Zheng, 2020; Flitsch et al., 2020). Quantifying these splice variants can reveal specific patterns associated with different cellular conditions, like neuronal differentiation, or disease states, like Alzheimer’s disease (AD) and other tauopathies (Hutton et al., 1998; J. Liu et al., 2018; Liu et al., 2022; Raj et al., 2018; Cieply & Carstens, 2015).

Although traditional short-read RNA sequencing (RNA-seq) has provided invaluable insights into the transcriptome, its limitations—such as difficulty in resolving complex transcript structures and accurately quantifying AS events—are well-documented (Conesa et al., 2016; Steijger et al., 2013; Trapnell et al., 2010). Short-read RNA-seq often struggles with repetitive regions and long contiguous sequences, leading to incomplete transcript assemblies (Bayega et al., 2018; Steijger et al., 2013; Tian et al., 2021). In contrast, LR sequencing technologies, such as those developed by Oxford Nanopore and PacBio, have revolutionized transcriptomics by producing reads that span entire transcripts, enabling accurate reconstruction of full-length RNA molecules (Amarasinghe et al., 2020; Oikonomopoulos et al., 2020). This breakthrough allows for the identification of novel transcripts and complex splicing events that are often missed by short-read methods (Aguzzoli Heberle et al., 2024; Inamo et al., 2024; Ulicevic et al., 2024; Wright et al., 2022). By mapping the full landscape of AS and transcript diversity, LR sequencing offers unprecedented precision in understanding gene regulation and its implications for human health and disease.

Analyzing differential transcript expression (DTE) provides critical insights into how different isoforms of the same gene are expressed under various conditions, offering a deeper understanding of gene regulation at the transcript level. Differential transcript usage (DTU), on the other hand, highlights shifts in the relative abundance of different transcript isoforms, indicating how splicing regulation may differ between conditions. For example, the *APP* gene, which encodes amyloid-beta precursor protein, and the presenilin 1 (*PSEN1*) gene undergo AS that directly influences the production of amyloid-beta peptides, a hallmark of AD pathology (Baulac et al., 2003; Cruts & Van Broeckhoven, 1998; Hutton, 1997; O’Brien & Wong, 2011). Moreover, *APP* and the bridging integrator 1 (*BIN1*) gene exhibit significant differential transcript expression and usage, especially in the temporal and frontal lobes of AD patients, emphasizing the pivotal role of AS in AD (Marques-Coelho et al., 2021).

In Parkinson’s disease (PD), several DTU events have been identified in the prefrontal cortex, with 23 specific events in 19 genes, including *THEM5, SLC16A1*, and *BCHE*, replicated across multiple individuals in PD cohorts, demonstrating the potential functional consequences of altered splicing (Dick et al., 2020). Similarly, in Huntington’s disease (HD), the *HTT* gene produces multiple isoforms through AS, which impact the aggregation propensity of the huntingtin protein, a central factor in HD etiology and progression (Sathasivam et al., 2013). As illustrated by these examples, investigating DTE and DTU during cell differentiation is essential for understanding the molecular basis of the homeostatic state, as well as neurodevelopmental and neurodegenerative diseases.

In our study, we utilized nanopore RNA-seq to achieve high-quality transcriptome profiling and a comprehensive analysis of DGE, DTE and DTU between human skin-derived fibroblasts, fibroblast-derived induced pluripotent stem cells (iPSCs), and iPSC-derived cortical neurons (iCNs). We identified key transcripts specific to each cell type and provided a detailed analysis of DTE between these cell types. Our DTU analysis revealed differences in transcript usage between cell types or during cell differentiation, likely driven by AS events. Notably, we identified genes with significant changes in transcript expression and usage, including medically relevant genes such as *APP*, *KIF2A*, and *BSCL2*, which undergo AS. These findings underscore the complexity of transcript regulation and highlight the importance of incorporating transcript-level analyses to fully understand transcriptomic regulation and disease mechanisms.

## Results

### Long Read Sequencing Uncovers Transcript Diversity and Complexity Across Cell Types

We utilized the Oxford Nanopore Technologies long-read sequencing platform to generate an average of 17,722,607 QC-passed reads, enabling us to assess transcript diversity across various cell types and differentiation stages, including fibroblasts, iPSC, and iCN. The properties and purity of cultured iCN were previously shown (Xu et al., 2024). Gene and transcript level quantification was performed using our robust pipeline (see methods). A transcript was considered expressed above noise levels if its median TPM exceeded one in each investigated cell type. Using this criterion, we identified a total of 21,040 unique transcripts across all three cell types with an average of 15,072 transcripts in iCN, 13,048 in fibroblasts, and 12,759 in iPSC (Figure 1A; Supplementary Figure S1). Additionally, our analysis revealed a significant number of lowly expressed transcripts (TPM between 0.1 and 1), with a count of 20,939 in iCN, 18,896 in fibroblasts, 19,571 in iPSC (Supplementary Figure S2A).

**Figure 1:**
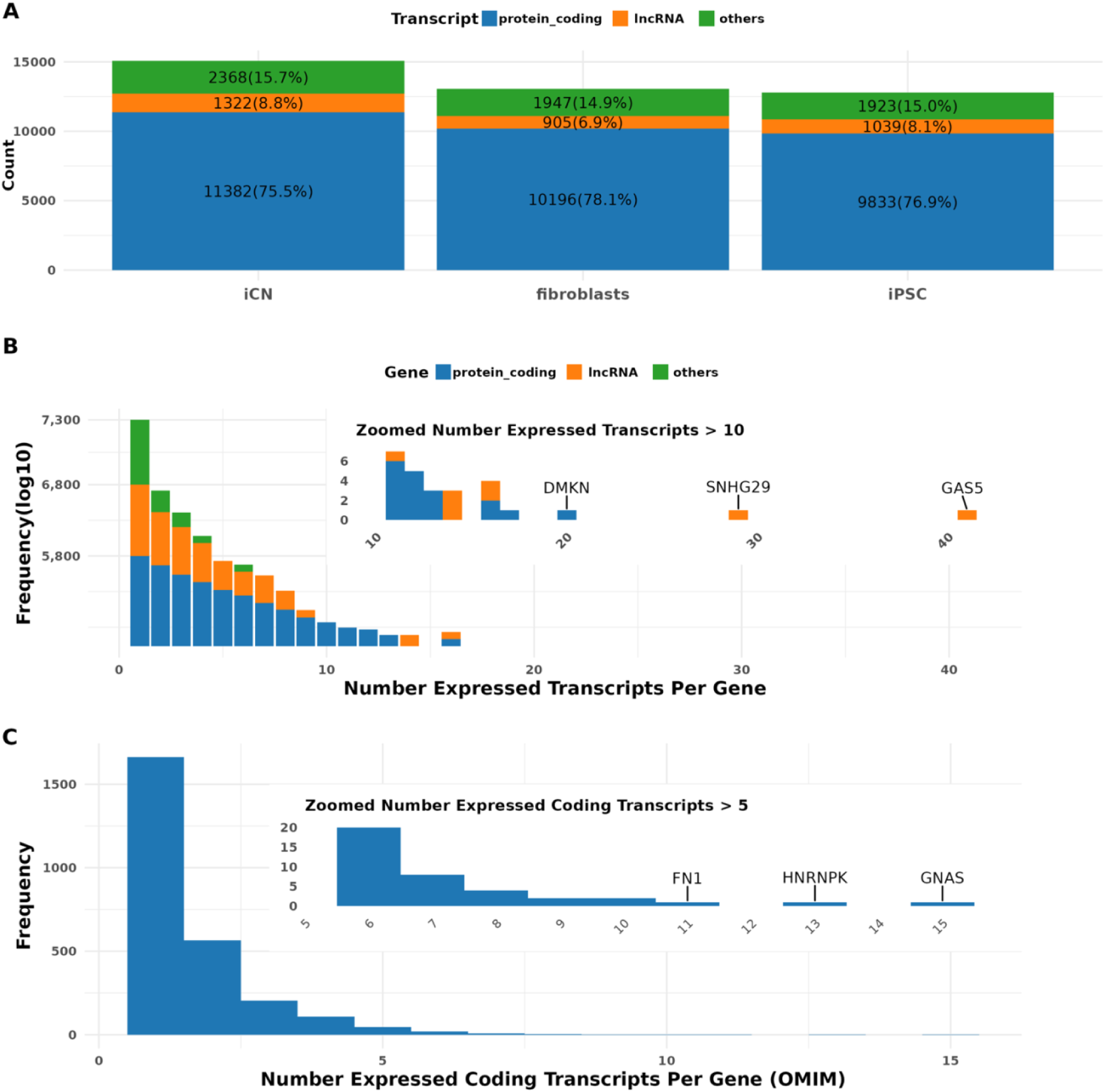
Transcriptome Profiling of Cultured Cells Using Long-Read RNA Sequencing: **(A) Transcript Type Distribution**: Bar plot showing counts and percentages of protein-coding transcripts (blue), long non-coding RNA (lncRNA) transcripts (orange), and other transcript types (green) in iCN (right), fibroblasts (middle), and iPSC (right**). (B) Transcript Diversity per Gene**: The frequency distribution of transcripts expressed per gene, categorized by biotype. The main plot shows the overall distribution, while the insert highlights genes with over 10 expressed transcripts, including DMKN, SNHG29, and GAS5. The bar length of each gene annotation is log10-transformed frequency and labeled with original frequency. **(C) Coding Transcript Frequency**: The distribution of the number of coding transcripts per gene for medically relevant genes as defined in the OMIM database. Most genes express fewer than 5 coding transcripts. The insert highlights genes with higher transcript variability, such as GNAS and HNRNPK.

To assess the accuracy of our LR transcriptomic data across cell types, we implemented a multi-step validation process. Initially, we curated a set of marker genes specific to neuron (n=85), fibroblasts (n=132) and iPSC (n=13) from the PanglaoDB database (https://panglaoDB.se/index.html). We examined the expression of these marker genes within our samples (Supplementary Figure S2B). The expression patterns of marker genes matched the expected cell types. Furthermore, we compared our results to previously published datasets, including two Illumina short-read datasets from Tsankov et al. (2015) and Burke et al. (2020) and a nanopore LR dataset from Ulicevic et al. (2024) (Burke et al., 2020; Tsankov et al., 2015; Ulicevic et al., 2024). Principal component analysis (PCA) demonstrated that our data clustered distinctly by cell type, showing clear separation and alignment with the existing datasets (Supplementary Figure S2C), further validating the accuracy of our data.

Transcripts identified in our samples were classified based on the Gencode annotation (v43) into categories such as protein-coding transcripts, long non-coding RNAs (lncRNAs), and other biotypes, including transcripts containing retained introns or subjected to nonsense mediated decay and pseudogene transcripts. On average, 76% of the expressed transcripts in our samples were protein-coding, 8% were lncRNAs, and 15% belonged to other biotypes across all investigated cell types. Specifically, in iCN, 11,382 (75.5%) transcripts were classified as protein-coding, 1,322 (8.8%) as lncRNAs, and 2,368 (15.7%) as other biotypes (Figure 1A).

We also investigated the number of unique transcripts observed per gene across cell types. Using our expression threshold, we detected 21,040 expressed transcripts from 11,853 unique genes. Interestingly, 60.8% (7,201) of these genes expressed only a single transcript (Figure 1B). On the other end of the spectrum, three genes demonstrated the expression of more than 20 transcripts each, including two lncRNA genes—*GAS5* (45 transcripts) and *SNHG29* (28 transcripts)—and one protein-coding gene, *DMKN* (20 transcripts) (Figure 1B). Additionally, we identified 5,135 coding transcripts from 2,740 genes defined as medically relevant according to the OMIM database (released at 2023.7) (Figure 1C). Among these, 1,207 (44.5%) genes expressed multiple transcripts (Figure 1C). Notably, genes such as the highly complex *GNAS*, which is involved in the assembly of the stimulatory G-protein alpha subunit and possibly involved in imprinting, the novel internal ribosomal entry site-transacting factor *HNRNPK*, and Fibronectin-1 (*FN1*), related to cell growth, differentiation and migration, exhibited over 10 transcripts each (Figure 1C).

### Differential Transcript Expression Reveals Key Transcripts Distinguishing Cell Types and Highlighting Disease Associations

Next, we utilized LR RNA-seq data to analyze differential transcript expression (DTE) across various cell types and identified 35,519 DTE events affecting 16,886 unique transcripts across 10,215 genes. Among these comparisons, 12,303 DTE events (Wald test with Benjamini-Hochberg (BH) correction, p < 0.05) were identified in the comparison of iCN vs. iPSC, 12,293 events (Wald test with BH correction, p < 0.05) in iCN vs. fibroblasts, and 10,923 events (Wald test with BH correction, p < 0.05) in fibroblasts vs. iPSC (Figure 2A). The analysis revealed a significant overlap of DTE across the comparisons. 4,262 DTE events were observed in all three comparisons. In contrast, a smaller subset of DTE transcripts was specific to individual comparisons. For example, 988 transcripts were unique to the iCN vs. iPSC comparison (Supplementary Figure S3A).

**Figure 2:**
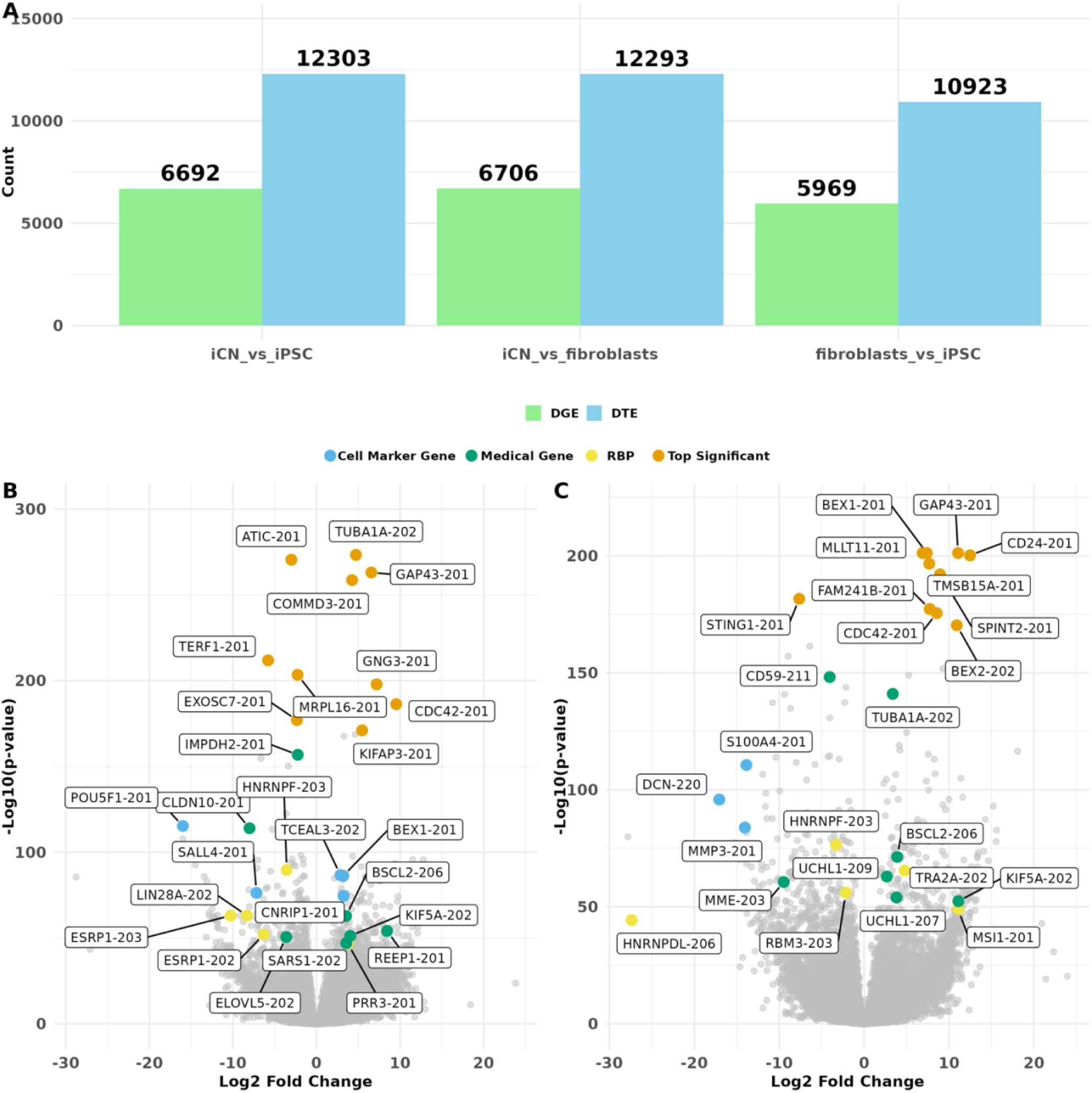
Differential Gene and Transcript Expression Analysis: **(A) Bar Plot of Differentially Expressed Genes (DGE) and Differentially Expressed Transcripts (DTE) Events**: Bar plot showing the number of DGEs (green) and DTEs (blue) identified in three comparisons: iCN vs. iPSC (left), iCN vs. fibroblasts (middle), and fibroblasts vs. iPSC (right). **(B) Volcano Plot for DTE of iCN vs. iPSC**: Volcano plot displaying the log2 fold change (x-axis) against the -log10 (adjusted p-value) (y-axis) for each transcript. A set of highlighted significant DTE events (adjusted p < 0.05 and |log2 fold change| > 1) are selected and categorized as top high-ranking events by p-value (orange), originating from medical-associated genes defined by OMIM (green), from RBP mRNAs (yellow), and from cell marker transcripts (blue). **(C) Volcano Plot for DTE of iCN vs. fibroblasts**: Volcano plot displaying log2 fold change (x-axis) against the -log10 (adjusted p-value) (y-axis) for each transcript. A set of highlighted significant DTE events (adjusted p < 0.05 and |log2 fold change| > 1) are selected and categorized as top high-ranking events by p-value (orange), originating from medical-associated genes defined by OMIM (green), from RBP mRNAs (yellow), and from cell marker transcripts (blue). Each comparison group included 9 independent replicates (n = 9)

Among DTE events identified in the iCN vs iPSC comparison, transcripts such as *TUBA1A-*202, a major component of microtubules crucial for cell structure and intracellular transport (Tantry & Santhakumar, 2023), *ATIC-*201, which plays an essential role in cellular proliferation (Niu et al., 2022), and *GAP43-* 201, known for its role in neuronal growth, regeneration, and synaptic plasticity (Chung et al., 2020; Holahan, 2015 ;M. R. Holahan, 2017; Kawasaki et al., 2018), exhibited the highest significance and fold change. Additionally, several transcripts from RNA-binding proteins (RBPs) (e.g., *HNRNPF-*203*, LIN28A-*202*, ESRP1-*202/203) were differentially expressed (Figure 2B). Disease-associated genes (e.g., *REEP1*-201*, KIF5A*-202*, EXOSC5*-201) were also identified, alongside the upregulation of neuronal marker gene transcripts (e.g., *BEX1*-201*, CNRIP1-*201*, TCEAL3-*202) and the expected downregulation of iPSC marker transcripts (e.g., *POU5F1*-201*, SALL4-*201) in iCN (Figure 2B).

Similarly, the comparison of iCN vs. fibroblasts revealed significant differential expression of transcripts (Figure 2C) such as *BEX1*-201, implicated in neurogenesis and neuronal differentiation (Jiang et al., 2016; Vilar et al., 2006), *GAP43-*201, plays a crucial role in nervous system development (Benowitz & Routtenberg, 1997), and *MLLT11-*201, involved in the development and progression of human cancers (Proestling et al., 2023). This comparison also highlighted transcripts linked to RNA-binding proteins (*e.g.*, *HNRNPF-*203, *RBM3*-203) and fibroblast marker genes (*e.g.*, *S100A4-*201*, CD24-*201) (Figure 2C). Furthermore, transcripts from medical or disease-related genes, such as *BSCL2-*206*, REEP1-*201*, UCHL1-*207/209 *and KIF5A-*202, all associated with Hereditary Spastic Paraplegia (HSP), were significantly differentially expressed. The comparison of fibroblasts vs. iPSC also highlighted key transcripts associated with cell marker genes (*e.g.*, *S100A4-*201*, POU5F1*-201), RBPs (*e.g.*, *LIN28A-* 202), and disease-related genes (*e.g.*, *MME-*203/214*, KIF1A-*207), which may play a role in maintaining fibroblast identity (Supplementary Figure S3B).

In addition to DTE, we investigated differential gene expression (DGE) to provide a broader context for our findings. Our analysis identified 19,367 DGE events (Wald test with BH correction, p < 0.05) across pairwise comparisons between cell types, involving 9,200 unique genes. Among these, the comparison of iCN vs. fibroblasts exhibited the most DGE events (n=6,706), followed by the comparison of iCN vs. iPSC (n=6,692) and fibroblasts vs. iPSC (n=5,969) (Figure 2A, Supplementary Figure S4A).

In the DGE analysis of iCN vs. iPSC, genes such as *TUBA1A, COMMD3, ATIC* and *GNG3* exhibited high significance and fold change, mirroring the patterns observed in the DTE analysis. Several other genes, including *REEP1, KIF5A, LIN28A* and *POU5F1*, were also prominent in both DGE and DTE analyses (Supplementary Figure S4B). Similarly, in the comparisons of iCN vs. fibroblasts and fibroblasts vs iPSC, several genes were consistently detected in both DGE and DTE analyses (Supplementary Figure S4C–D). However, some genes, such as *SGCE* and *STRADA* (Table 1), did not show significant differential expression at the gene level between cell types but exhibited distinct differential expression patterns at the transcript level (Supplementary Figure S7A, B).

**Table 1:**
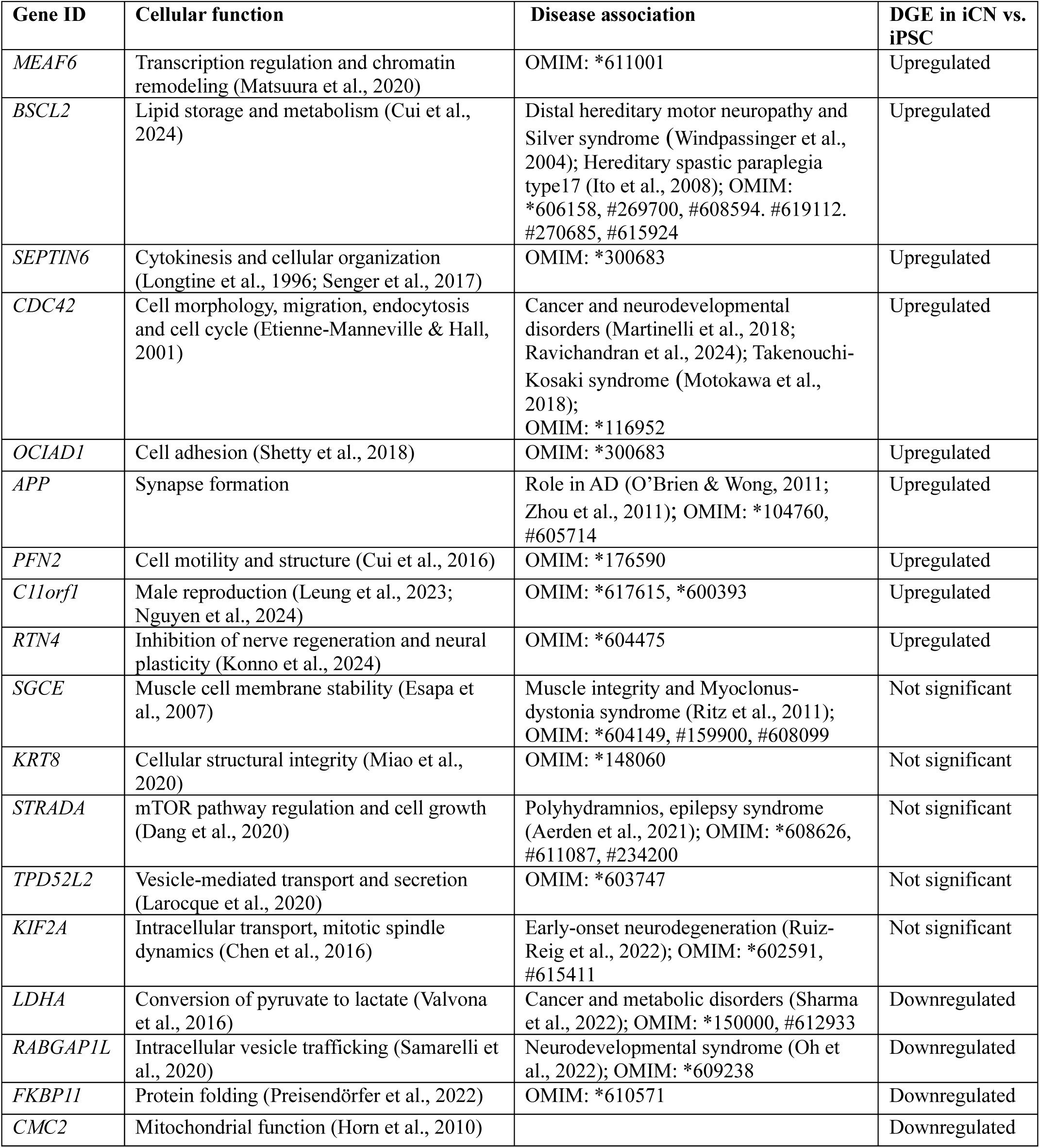
Summary of significant DTU genes between iCN and iPSC. The table lists key genes, their cellular function, disease association, and their regulation status at gene level (upregulated, downregulated or unchanged) in iCN from our DGE analysis.

| Gene ID | Cellular function | Disease association | DGE in iCN vs. iPSC |
| --- | --- | --- | --- |
| *MEAF6* | Transcription regulation and chromatin remodeling (Matsuura et al., 2020) | OMIM: *611001 | Upregulated |
| *BSCL2* | Lipid storage and metabolism (Cui et al., 2024) | Distal hereditary motor neuropathy and Silver syndrome (Windpassinger et al., 2004); Hereditary spastic paraplegia type17 (Ito et al., 2008); OMIM: *606158, #269700, #608594. #619112. #270685, #615924 | Upregulated |
| *SEPTIN6* | Cytokinesis and cellular organization (Longtine et al., 1996; Senger et al., 2017) | OMIM: *300683 | Upregulated |
| *CDC42* | Cell morphology, migration, endocytosis and cell cycle (Etienne-Manneville & Hall, 2001) | Cancer and neurodevelopmental disorders (Martinelli et al., 2018; Ravichandran et al., 2024); Takenouchi-Kosaki syndrome (Motokawa et al., 2018); OMIM: *116952 | Upregulated |
| *OCIAD1* | Cell adhesion (Shetty et al., 2018) | OMIM: *300683 | Upregulated |
| *APP* | Synapse formation | Role in AD (O'Brien & Wong, 2011; Zhou et al., 2011); OMIM: *104760, #605714 | Upregulated |
| *PFN2* | Cell motility and structure (Cui et al., 2016) | OMIM: *176590 | Upregulated |
| *C11orf1* | Male reproduction (Leung et al., 2023; Nguyen et al., 2024) | OMIM: *617615, *600393 | Upregulated |
| *RTN4* | Inhibition of nerve regeneration and neural plasticity (Konno et al., 2024) | OMIM: *604475 | Upregulated |
| *SGCE* | Muscle cell membrane stability (Esapa et al., 2007) | Muscle integrity and Myoclonus-dystonia syndrome (Ritz et al., 2011); OMIM: *604149, #159900, #608099 | Not significant |
| *KRT8* | Cellular structural integrity (Miao et al., 2020) | OMIM: *148060 | Not significant |
| *STRADA* | mTOR pathway regulation and cell growth (Dang et al., 2020) | Polyhydramnios, epilepsy syndrome (Aerden et al., 2021); OMIM: *608626, #611087, #234200 | Not significant |
| *TPD52L2* | Vesicle-mediated transport and secretion (Larocque et al., 2020) | OMIM: *603747 | Not significant |
| *KIF2A* | Intracellular transport, mitotic spindle dynamics (Chen et al., 2016) | Early-onset neurodegeneration (Ruiz-Reig et al., 2022); OMIM: *602591, #615411 | Not significant |
| *LDHA* | Conversion of pyruvate to lactate (Valvona et al., 2016) | Cancer and metabolic disorders (Sharma et al., 2022); OMIM: *150000, #612933 | Downregulated |
| *RABGAP1L* | Intracellular vesicle trafficking (Samarelli et al., 2020) | Neurodevelopmental syndrome (Oh et al., 2022); OMIM: *609238 | Downregulated |
| *FKBP11* | Protein folding (Preisendörfer et al., 2022) | OMIM: *610571 | Downregulated |
| *CMC2* | Mitochondrial function (Horn et al., 2010) |  | Downregulated |

### Differential Transcript Usage Highlights Transcript Complexity and Functional Adaptations Across Cell Types

Differential transcript usage (DTU) refers to variations in the relative abundance of transcripts from the same gene across conditions or cell types. These variations often indicate functional adaptations at the transcript level. Genes were considered to exhibit DTU if they had at least one significant DTU event. Comparisons between cell types revealed a total of 5,135 DTU events involving 3,851 unique transcripts from 1,894 unique genes. These were observed among 4,652 genes that expressed more than one transcript in our dataset. The highest number of DTU events was observed in the comparison of iCN vs. iPSC (n=2,109), followed by fibroblasts vs. iPSC (n=1,651), and iCN vs. fibroblasts (n=1,375) (Figure 3A). 1,332 DTU events were unique to the iCN vs. iPSC comparison, indicating distinct transcript usage patterns in this transition (Figure 3A). Interestingly, we observed 332 shared DTU events between all comparisons, which might suggest common regulatory mechanisms (Figure 3A).

**Figure 3:**
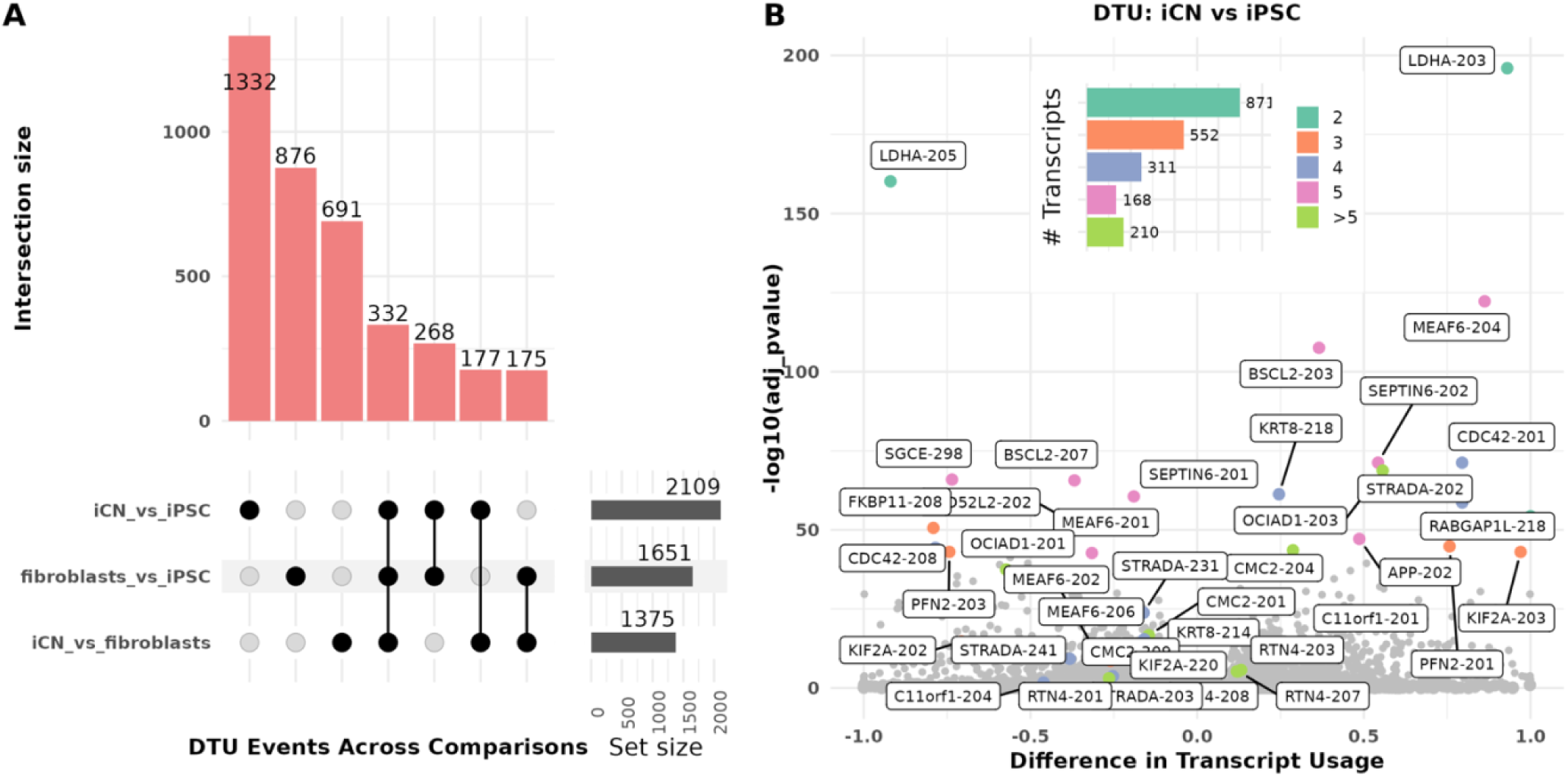
Differential Transcript Usage (DTU) Analysis Across Cell Types. **(A) UpSet Plot**: UpSet plot illustrating the intersection of DTU events across three comparisons: iCN vs. iPSC, fibroblasts vs. iCN, and fibroblasts vs. iPSC. The bars indicate the number of shared and unique DTU transcripts for each comparison. iPSC vs. iCN comparison has the highest number of unique DTU transcripts (1,332). The bar plot denotes the intersection size, circles denote which comparisons have overlap, and the set size reflects the total number of genes **(B) Volcano Plot for iCN vs. iPSC Comparison:** Volcano plot showing the difference in transcript usage (x-axis) against the -log10 (adjusted p- value) (y-axis) for each transcript. Top significant DTUs are highlighted, with notable genes such as APP, KIF2A and BSCL2. Points are colored based on the number of transcripts per gene involved in the DTU event, as indicated by the insert bar chart (aquamarine = 2 transcripts (n=871); orange= 3 transcripts (n=552); purple = 4 transcripts (n=311); pink = 5 transcripts (n=168); green > 5 transcripts (n=210)). The highest frequency of DTU events occurs in genes with two expressed transcripts, followed by those with three and four transcripts. Each comparison group included 9 independent replicates (n = 9).

In the comparison between iCN and iPSC, the top-ranking DTU events, based on statistical significance, revealed several significant findings. These include genes involved in essential cellular functions such as metabolism, cytoskeletal organization, and vesicle trafficking. Table 1 summarizes 18 top ranking DTU genes, their cellular function and disease association, along with gene-level expression (DGE) changes (upregulated, downregulated or not significant in iCN vs. iPSC).

Analyzing significant DTU events in this comparison revealed that genes with two expressed transcripts were the most common (n=871) (Figure 3B). A similar pattern was observed in the comparison of iCN vs. fibroblasts (n=566) and fibroblasts vs. iPSC (n=689) (Supplementary Figure S5A-D). Additionally, many genes exhibited DTU events with more than two expressed transcripts (Figure 3B). For example, in the iCN vs. iPSC comparison, genes such as *RTN4, BSCL2,* and *APP* showed multiple transcript variants, highlighting the high level of transcript complexity in these genes (Figure 3B).

*CDC42* displayed differential expression and transcript usage at both the gene and transcript levels (Supplementary Figure S6A). In iCN, *CDC42* was upregulated at the gene level, with transcript *CDC42*- 201 predominantly expressed and showing higher usage (Supplementary Figure S6B). In contrast, *CDC42*-208 was the transcript predominantly used in iPSC (Supplementary Figure S6B). *LDHA* was downregulated at the gene level in iCN but exhibited a complex transcript regulation pattern, with some transcripts upregulated and others downregulated, alongside differential transcript usage between iCN and iPSC (Supplementary Figure S6B). Interestingly, other medically relevant DTU genes, such as *SGCE* and *STRADA*, did not show statistically significant changes at the gene level (Supplementary Figure S7A, B). However, both genes exhibited multiple differentially expressed transcripts and notable shifts in transcript usage. For instance, four *STRADA* transcripts (*STRADA-*202*, STRADA-*203*, STRADA-*231 and *STRADA-*241) were identified in our dataset (Supplementary Figure S7B). *STRADA-*202 showed the highest transcript abundance and usage in iCN, while the other three transcripts exhibited higher expression levels and greater usage frequency in iPSC (Supplementary Figure S7B).

To explore the relationship between differential regulation at the gene and transcript levels (DTE, DTU, and DGE) in greater detail, we focused on the medically relevant DTU genes *APP*, *KIF2A* and *BSCL2*. It is widely accepted that *APP* is an Alzheimer hallmark gene and plays a significant role as a regulator in neural system development (O’Brien & Wong, 2011). The alternative splicing of *APP* mRNA can generate approximately ten different transcripts, which are differentially expressed in various tissues (Coronel et al., 2019; Jakubauskienė & Kanopka, 2021). In our analysis, we observed upregulation of *APP* in iCN at gene level (DGE) and identified five distinct *APP* mRNA transcripts (Figure 4A). Of these, four (*APP-*201*, APP-*202*, APP-*204, and *APP-*205) are annotated as protein-coding, while one, *APP-*218, is classified as protein-coding, though its coding sequence (CDS) has not been fully defined in the latest Gencode annotation (Figure 4A). Our differential transcript analysis revealed that three transcripts (*APP-* 201*, APP-*202, and *APP-*218) displayed differential expression and usage in iCN compared to iPSC (Figure 4A). Interestingly, the transcript *APP-*202, also known as *APP695*, which lacks the Kunitz-like protease inhibitor (KPI) domain and Ox-2 antigen domain, was predominantly expressed in iCN, while *APP-*201 *(APP770)* was primarily expressed in iPSC (Figure 4A).

**Figure 4:**
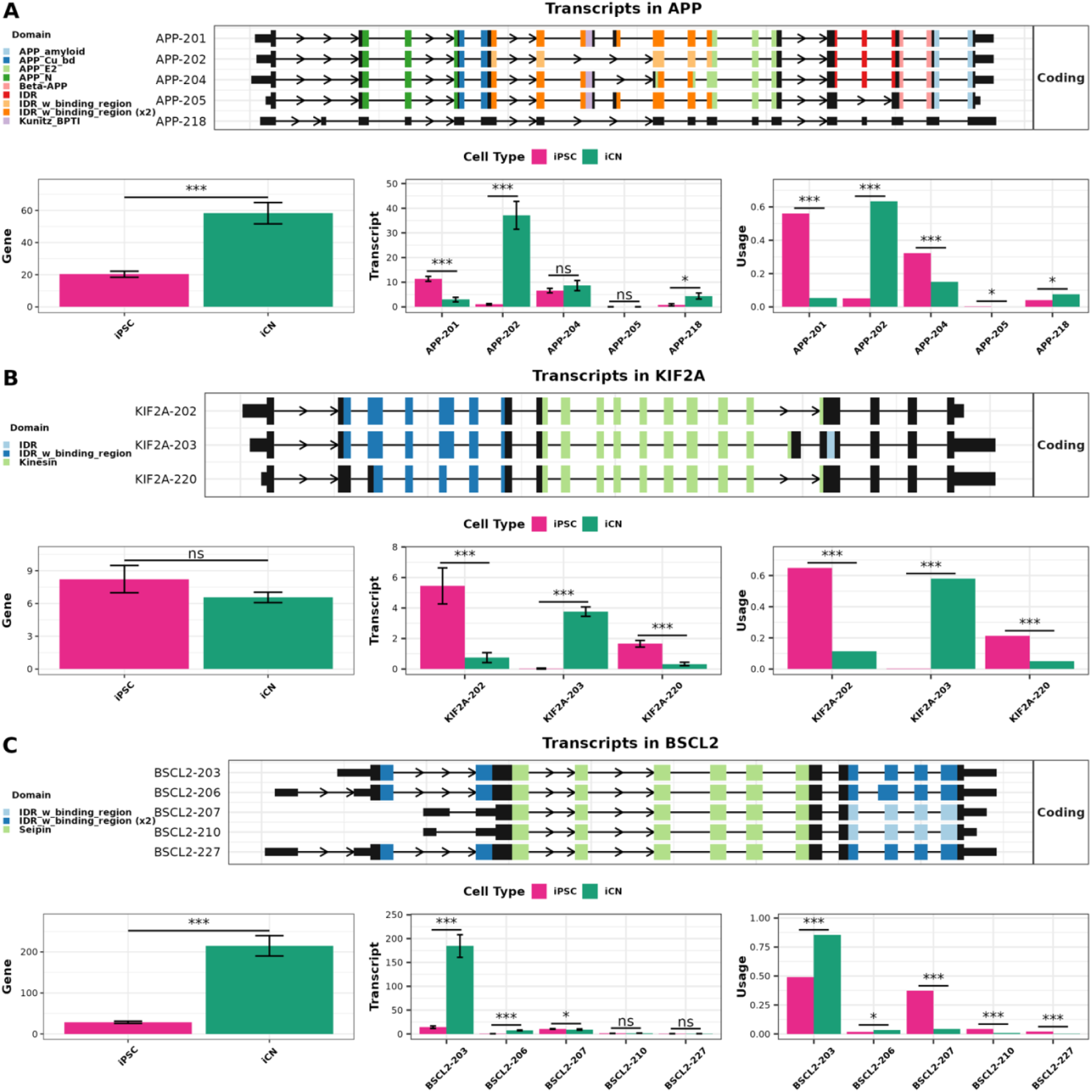
Differential Transcript Usage Analysis for the APP, KIF2A and BSCL2 Genes. For each of the selected genes **(A)** APP Gene; **(B)** KIF2A Gene; **(C)** BSCL2 Gene, the top panel indicates the transcript structures based on Gencode annotation v43, with main protein domains indicated by different colors. Bottom Panels: Gene expression (normalized TPM) levels (left), transcript expression (normalized TPM) levels (middle), and transcript usage (right) in iPSC (pink) and iCN (green). Statistical significance is indicated by ns (not significant), * (p < 0.05), and *** (p < 0.001). Each comparison group included six independent replicates (n = 9).

*KIF2A (*Kinesin Superfamily Protein 2A) plays a crucial role in neuronal migration and differentiation, primarily through its involvement in microtubule dynamics. Mutations in *KIF2A* are associated with cortical dysplasia and early-onset neurodegeneration (Ruiz-Reig et al., 2022; Homma et al., 2003). Alternative splicing of *KIF2A* mRNA produces multiple isoforms, each with distinct functional roles. Specifically, in mice, the inclusion or exclusion of exon 18, along with the alternative 5′ splice site selection in exon 5, generates isoforms that differ in their ability to support neuronal migration (Akkaya et al., 2021). In our study, three *KIF2A* transcripts (*KIF2A-*202*, KIF2A-*203, and *KIF2A-*220) were identified, with significant differential expression and usage between iCN and iPSC (Figure 4B). Notably, *KIF2A-*202 and *KIF2A-*220 were predominantly expressed in iPSC, while *KIF2A-*203, an isoform implicated in neuronal migration (Akkaya et al., 2021), was more prevalent in iCN (Figure 4B).

Importantly, despite the differential expression and usage of these transcripts, the overall gene expression of *KIF2A* remained unchanged between iPSC and iCN (Figure 4B). This finding suggests that while *KIF2A* is subject to complex regulation at the transcript level, its overall gene expression is tightly controlled, possibly to maintain essential cellular functions.

*BSCL2* (Berardinelli-Seip Congenital Lipodystrophy 2) encodes Seipin, an integral endoplasmic reticulum membrane protein crucial for lipid droplet formation and metabolism. Mutations in *BSCL2* are associated with congenital generalized lipodystrophy type 2, as well as neurodegenerative axonopathies such as hereditary spastic paraplegia and distal hereditary motor neuropathy (Hsiao et al., 2016; Li et al., 2022; Sánchez-Iglesias et al., 2021). Aberrant splicing of *BSCL2* has been implicated in several pathological conditions (Guillén-Navarro et al., 2013). In our analysis, we detected several *BSCL2* transcripts (*BSCL2*-203, *BSCL2*-206, *BSCL2*-207, *BSCL2*-210, and *BSCL2*-227) with differential expression and usage in iCN compared to iPSC (Figure 4C). Notably, *BSCL2*-203 was predominantly expressed in iCN, while *BSCL2*-207 was more prevalent in iPSC, though expressed at significantly lower levels (Figure 4C).

### Comparative Analysis of DGE, DTE, and DTU Reveals Overlapping and Unique Gene Functional Patterns

We conducted a direct comparison of the genes identified in each analysis (DGE, DTE, and DTU). The greatest overlap was observed between DGE and DTE genes, with a smaller proportion of significant genes identified by all three methods (Figure 5A-C).

**Figure 5:**
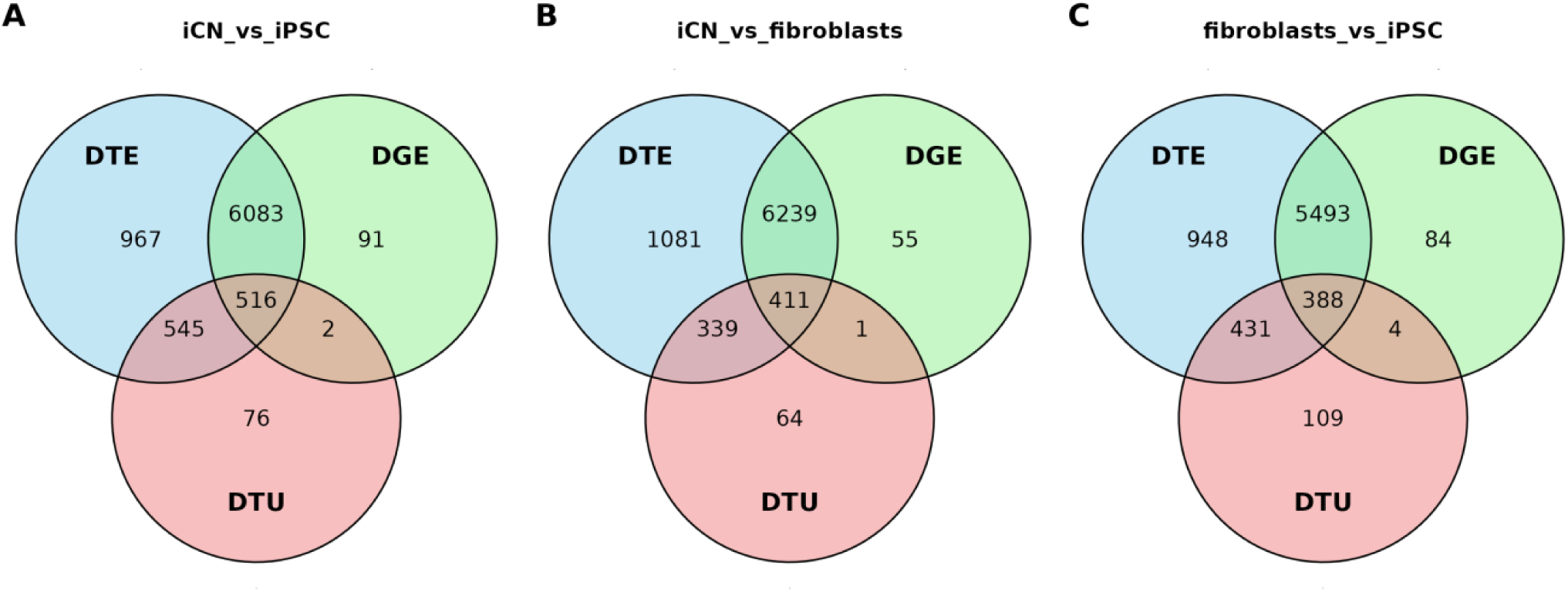
Venn Diagrams of Differential Gene Expression (DGE), Differential Transcript Expression (DTE), and Differential Transcript Usage (DTU) Across Cell Type Comparisons. Venn diagrams illustrating the overlap between DGE, DTE, and DTU genes in three different cell type comparisons: **(A)** iCN vs. iPSC, **(B)** iCN vs. fibroblasts, **(C)** fibroblasts vs. iPSC. Each circle represents one of the categories (DTE: top right; blue, DGE: top left; green; DTU: bottom; red), with the numbers indicating the count of genes in each category and their intersections.

In the comparison of iCN vs. iPSC, 6,559 genes were differentially expressed both at the gene and transcript level, with 516 genes shared across all three differential analyses (Figure 5A). Similar patterns were observed in the comparisons of iCN vs. fibroblasts (Figure 5B) and fibroblasts vs. iPSC (Figure 5C). Interestingly, a notable proportion of DTE genes were not found in the DGE analysis, suggesting that transcript-level regulation plays a pivotal role in cell-specific functions, which may be overlooked by solely investigating gene-level expression. The DTU analysis identified the smallest set of genes, many of which overlap with either DGE or DTE, suggesting that changes in transcript usage often coincide with changes in overall transcript or gene expression levels. The small number of unique DTU genes indicates that few genes exhibit changes in transcript usage without showing changes in transcript abundance or gene expression.

We performed functional enrichment analysis using gprofiler2 on DGE, DTE, and DTU genes across all comparisons. The top 10 GO terms with the best adjusted p-values of enrichments from each category— biological processes (BP), cellular components (CC), and molecular functions (MF)—are displayed in Figure 6 and Supplementary Figures S8 and S9. In the iCN vs. iPSC comparison, we found significant enrichment of 819 (DGE), 882 (DTE), and 169 (DTU) GO terms (BH-adjusted p-value < 0.05). Many of these terms were related to neuronal functions, with “neuron,” “synapse,” and “axon” prominently appearing in DGE (150 terms, 18%) and DTE (137 terms, 15%), reflecting biological processes related to cellular regulation, neurogenesis, and neuron development (Figure 6A). Additionally, both DGE and DTE genes in the iCN vs. iPSC comparison were enriched for cellular components of intracellular anatomical structures and organelles (Figure 6B). While DGE and DTE showed significant neuronal enrichment, the DTU gene set demonstrated a different pattern. Here, neuronal pathways were less enriched, with only one significant term, GO:0048699 (generation of neurons) (Figure 6A). Instead, DTU genes were more prominently associated with molecular functions related to cadherin and DNA binding (Figure 6C), pointing to additional transcriptomic regulatory layers that may not be fully captured by DGE and DTE analyses alone. Altogether similar pathways were enriched in the comparison of iCN vs fibroblasts, indicating that the underlying regulatory profile is indeed driven by the distinct neuronal properties of the iCN. Details on GO enrichment analysis in iCN vs fibroblasts and fibroblasts vs iPSCs can be found in Supplementary Figures S8 and S9.

**Figure 6:**
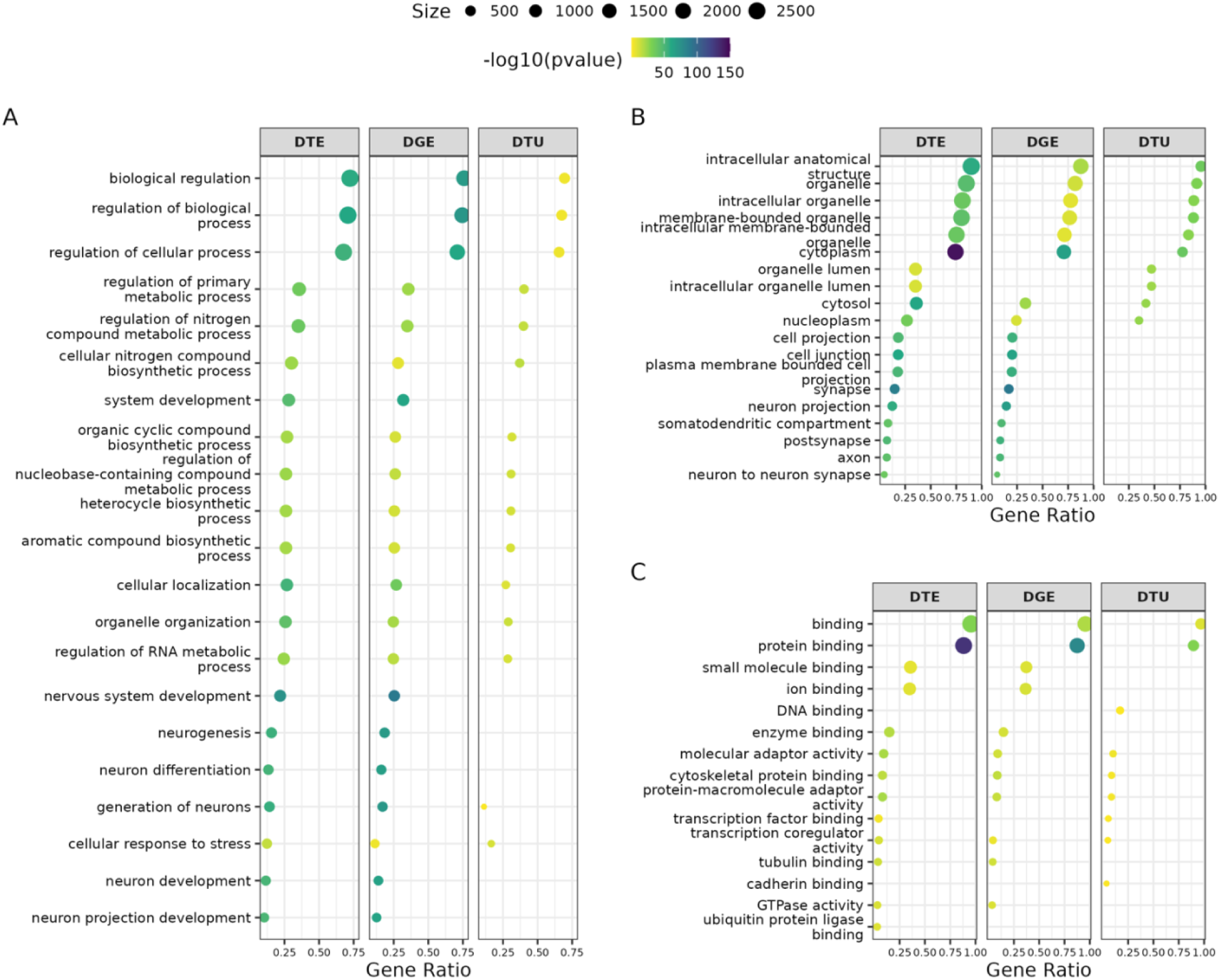
Gene Ontology (GO) Term Enrichment Analysis for DGE, DTE, and DTU in iCN vs iPSC. Dot plots illustrate the top 10 terms with the best adjusted p-values for enrichment from three categories: (**A**) biological processes (BP), **(B)** cellular components (CC), **(C)** and molecular functions (MF) for genes identified through DGE, DTE, and DTU analyses. Each dot represents a specific GO term, with the size of the dot indicating the number of associated genes and the color reflecting the -log10(p-value), signifying the level of statistical significance.

In conclusion, the cross-analysis of DGE, DTE, and DTU underscores the intricate layers of gene regulation at both the gene and transcript levels. The substantial overlap between DGE and DTE highlights their shared biological relevance, while the unique contributions from DTU emphasize the importance of considering transcript usage.

## Discussion

Our comprehensive transcriptome analysis using nanopore LR RNA-seq has provided valuable insights into the transcript diversity and expression profiles of human iPSC-derived iCN, fibroblasts, and iPSC. By integrating differential transcript expression (DTE) and differential transcript usage (DTU) analyses, we were able to uncover a level of transcriptomic complexity that extends beyond conventional gene expression studies. This approach has proven critical for identifying transcript-specific regulation and alternative splicing (AS) events, offering a more nuanced understanding of the functional implications of transcript variation between cell types.

The primary goal of our study was to characterize transcript expression and its changes during cellular differentiation. Our results revealed that iCN exhibited the highest number of expressed transcripts compared to fibroblasts and iPSC. This aligns with findings from Heberle et al. (2023) and Page et al. (2024), who also emphasized the expansive transcriptomic landscape required for proper neuronal function (Heberle et al. 2023; Page et al. 2024). This observation is consistent with the complex roles of neurons, which rely on a diverse set of transcripts to support critical processes like synaptic plasticity, neurotransmission, and neurogenesis.

Our study further underscores the power of DTE and DTU analyses in revealing the functional consequences of AS. For instance, our investigation into the amyloid precursor protein (*APP*) highlighted distinct splicing patterns that are relevant to Alzheimer’s disease (AD) (Jakubauskienė & Kanopka, 2021). Specifically, our combination of DTE and DTU analysis revealed that the neuron-specific isoform *APP695* (*APP-*202) was predominantly expressed in iCN, whereas *APP770 (APP-*201) and *APP751 (APP-*204) were more prevalent in iPSC. These findings align with the well-established role of *APP* in AD pathology, where *APP695*, lacking the KPI domain, is more susceptible to proteolytic cleavage, leading to the generation of amyloid-β peptides, a hallmark mechanism of AD (Zhang et al., 2011). The higher expression of KPI-containing isoforms in non-neuronal cells might suggest a protective mechanism against amyloidogenic processing (Belyaev et al., 2010; Golde et al., 1990; Hung et al., 1992; J. Kang et al., 1987; Weidemann et al., 1989). This observed pattern of isoform expression supports the validity of our transcript-level analyses and highlights the significance of transcript-specific regulation in understanding disease mechanisms.

Moreover, our study demonstrated that transcript differential analyses can uncover cell-type-specific transcript patterns that would otherwise be missed with gene-level analyses alone. For example, we identified key transcript isoforms in *KIF2A* that exhibited distinct splicing patterns during the transition from iPSC to iCN, even though overall gene expression levels showed no significant differences. Specifically, isoforms *KIF2A-*202 and *KIF2A-*220, which lack exon 17, were significantly upregulated according to DTE analysis and predominantly expressed in iPSC according to DTU analysis. In contrast, *KIF2A-*203, which retains exon 17, was upregulated in iCN based on DTE and showed predominant usage according to DTU analysis. This suggests that neurons utilize specific *KIF2A* isoforms during differentiation, likely to support microtubule depolymerization, a process critical for neuronal migration and axonal development (Miyamoto et al., 2015; Ruiz-Reig et al., 2022).

Importantly, transcript differential analyses (DTE and DTU) not only identify these transcript patterns but also guides formation of hypotheses on the regulatory mechanisms of AS. For example, the regulation of AS in *KIF2A* may be influenced by neuron-specific RNA-binding proteins (RBPs), which are known to bind to intronic sequences and 3’ UTRs, influencing exon inclusion and mRNA stability (Ince-Dunn et al., 2012; Lee et al., 2021). This mechanism could ensure the inclusion of exon 17 in *KIF2A-*203, preserving its role in microtubule depolymerization, essential for neuronal differentiation and migration (Akkaya et al., 2021). Conversely, isoforms lacking exon 17, such as *KIF2A-*202 and *KIF2A-*220, may serve broader, more generalized cellular roles.

One of the most significant contributions of DTE and DTU analyses is their potential to identify disease-causing targets. In the past, genomic mutations were often linked only to gene-level changes, providing limited insight into how these mutations impact specific transcript isoforms. However, our findings demonstrate that while a gene may exhibit straightforward regulation at the gene level, its transcripts can display complex and varied regulatory patterns, as exemplified by *APP* and *KIF2A*. This indicates that transcript-level changes, such as AS or DTU, can lead to the production of isoforms with distinct functional properties, even when gene expression remains unchanged. These transcript-specific variations may drive disease pathology by altering key cellular processes like protein function, localization, or molecular interactions. Therefore, transcript-level analyses allow us to directly link genomic mutations to specific transcript isoforms, offering more precise therapeutic targets for intervention.

For example, mutations in the *BSCL2* gene, such as the homozygous c.974dupG mutation, have been shown to lead to exon 7 skipping, potentially producing an abnormal transcript (Sánchez-Iglesias et al., 2021). This splicing alteration, which results in the loss of exon 7 in certain isoforms, has been linked to severe neurodegenerative conditions like Celia’s encephalopathy (Sánchez-Iglesias et al., 2021). In our study, we found that the *BSCL2-*203 transcript, which includes exon 7, is the dominant isoform of the Seipin family expressed in iCN. The presence of exon 7 in *BSCL2-*203 may be critical for maintaining normal neuronal function, and any disruption in this splicing pattern, such as the exon 7 skipping caused by the c.974dupG mutation, could lead to pathogenic outcomes. This example illustrates how transcript-specific changes, driven by AS mutations, can be directly tied to disease mechanisms, emphasizing the importance of identifying specific transcripts as potential therapeutic targets.

In conclusion, by leveraging nanopore LR RNA-seq, we were able to explore transcript diversity at a high resolution, providing a comprehensive framework for understanding isoform-specific regulation and its impact on cellular identity. This understanding is critical for advancing research into neurodevelopment and neurodegeneration. Importantly, we show that LR RNA-seq, combined with DTE and DTU analyses, can uncover the complexities of transcript regulation, particularly how AS shapes cellular identity and function.

Transcript-level analysis may prove crucial in the future, as genetic testing alone overlooks changes in isoform expression and AS that contribute to disease pathology. AS or alterations to the predominantly expressed isoforms in disease-relevant tissues may also account for diseases that currently lack a genetic explanation. Furthermore, identification of disease-relevant transcripts may open potential therapeutic avenues, as interventions targeting specific isoforms could restore normal splicing patterns and prevent the production of pathogenic isoforms (Nikom & Zheng, 2023). Ultimately, transcript-level analyses have the potential to significantly enhance our approach to genetic testing by identifying mutations that lead to aberrant splicing or isoform production. Taken together, these analyses hold significant promise for advancing our understanding of cellular functions, especially in neurodevelopmental and neurodegenerative diseases, where transcriptomic diversity and splicing dysregulation play pivotal roles. Moving forward, a continued focus on transcript-level changes in various disease models will provide deeper insights into the molecular mechanisms of disease progression and may lead to the identification of novel therapeutic targets for intervention.

## Methods

### Generation of human cell lines

The study was approved by the Institutional Review Board of the University of Tübingen Medical Faculty at the University Hospital Tübingen, Germany (IRB: 423/2019BO1). All participants gave their written informed consent to study participation. Human skin fibroblasts were obtained from healthy donors as previously described (Vangipuram et al., 2013). Fibroblasts were cultured in DMEM high glucose media (Sigma) with 10% FCS (Life Technologies). From these fibroblasts, iPSCs were reprogrammed as previously described (Manibarathi et al., 2024; Nagel et al., 2020), using episomal plasmids (pCXLE-hUL ID: #270776, pCXLE-hSK, ID: #27078, and pCXLE-hOCT3/4, ID: #27076 (Okita et al., 2011). The reprogrammed cells were seeded onto Matrigel-coated (1:60 in DMEM, Corning®) 6-well plates in fibroblast media. The next day, the media was supplemented with 2 ng/ml FGF2 (Peprotech) and 1% P/S (Gibco). On day three media was changed to Essential 8 (E8; in-house) media with 100 µM Sodium butyrate (Sigma-Aldrich) and 0.1% P/S and changed every other day. After 3 to 4 weeks, colonies were picked and expanded. Cryo-stocks were obtained using 50% E8 media with 40% KO-SR (Life Technologies), 10% DMSO (Sigma-Aldrich) and 10 μM Y-27632 (Abcam Biochemicals). A PCR Mycoplasma Test (AppliChem) was performed following manufacturer’s recommendation. Reprogrammed iPSCs were differentiated into cortical neurons of layers V and VI. The differentiation followed previously established protocols (Hauser et al., 2020; Schuster et al., 2020; Xu et al., 2024). Briefly, iPSCs were seeded at a density of 3 x 10^5^ cells/cm^2^ on Matrigel-coated plates (Corning®), in E8 medium supplemented with 10µM SB431542 (Sigma-Aldrich) and 500nM LDN- 193189 (Sigma-Aldrich). The cells underwent neural induction over 9 days. Post induction, on Day 9, the cells were split at a 1:3 ratio and then cultured in 3N medium with 20ng/ml FGF-2 for an additional 2 days. From Day 11 to Day 26 after induction (DAI 11-26), cells were maintained in 3N medium, with addition of heparin (100µg/ml) on DAI 13 and 15. Medium changes occurred bi-daily. iCNs were split on DAI 26, with supplementation of 10 µM DAPT (Tocris) and 10 µM PD0325901 (Tocris) in DAI 27 and 29. iCNs were maintained until DAI37 and RNA was isolated according to the manufacturer instructions (Qiagen; RNeasy Mini Kit). Cortical neuron identity was previously shown by immunocytochemistry (neuronal markers: β-III-tubulin, TAU; dendritic marker: MAP2; Cortical layer V (CTIP2) and VI (TBR1) markers) and RT-qPCR (Cortical layer markers: FOXG1 and PAX6; dendritic marker: MAP2; microtubule-associated marker: DXC) at DAI36 (Xu et al., 2024). A total of four cell lines were used for the experiments (Table 2). All cells described were maintained at 37 °C and 5 % CO2.

**Table 2:** Cell lines used for long-read RNA-sequencing experiments.

| Human-derived fibroblasts | Fibroblast-derived induced pluripotent stem cells | iPSC-derived cortical neurons |
| --- | --- | --- |
| CO-1 | CO-1 2.1 | CO-1 2.1 |
| - | CO-2 1.1 | CO-2 1.1 |
| CO-3 | CO-3 B.1 | CO-3 B.1 |
| CO-4 | - | - |

### Long-read RNA-sequencing processing and quality control

RNA concentration was estimated using the Qubit Fluorometric Quantitation and RNA Broad-Range Assay (Thermo Fisher Scientific) and RNA Integrity Number RIN using the Fragment Analyzer 5300 and the Fragment Analyzer RNA kit (Agilent Technologies). Libraries were prepared using the PCR-cDNA Barcoding Kit (SQK-PCB111.24) kit from Oxford Nanopore Technologies according to manufacturer’s instructions. A total of 200 ng of total RNA was annealed for strand-switching reaction and reverse transcription with Maxima H Minus RT (Thermo Fisher Scientific). The resulting cDNA was amplified with LongAmp Hot Start Taq Master Mix (NEB) according to the protocol from ONT; with following modifications; extension for 9 min with 12 cycles of amplification and a final extension of 10 minutes. Library molarity was determined by measuring the library size using the Femto*Pulse* instrument and the Genomic DNA 165 kb Kit (Agilent Technologies) and the library concentration using Qubit Fluorometric Quantitation and dsDNA High sensitivity assay (Thermo Fisher Scientific). An equimolar pool of 4 barcoded libraries was cleaned up with 0.8x AMPure XP beads (Beckman Coulter). The pools were quantified and assessed with Qubit and Femto, as individual libraries. The Rapid Adaptors were added to the pool of amplified cDNA and 20 fmol of the library was loaded on a R.9.4.1 PromethION flow cell (FLO-PRO002) and ran on a PromethION instrument in High Accuracy Basecalling mode for 72 hours.

Demultiplexing was performed by MinKNOW software. Initially, quality was assessed using the pycoQC tool (Leger et al., 2019). Reads were processed with Pychopper (https://github.com/nanoporetech/pychopper). The FASTQ files were mapped to the reference human genome (GRCh38) using the minimap2 tool with the splicing option and quantified using Bambu v3.2.4 (Y. Chen et al., 2023) to obtain transcript-level counts and CPM. Then CPM were converted to TPM (Transcripts Per Million) with Gencode annotation v43. TPM was used as expression values to select transcripts for a given cell type. Median values of transcript expression were then calculated per cell type and transcripts with median values less than a threshold 1 were filtered out for a given cell type. Finally, all transcripts expressed in each cell type were combined. Gene expression was then aggregated from transcript expression by summing all expressed transcripts in a gene.

Principal component analysis (PCA) was performed on three published datasets, along with our own dataset, using the *prcomp* function in R version 4.3.1. Prior to PCA, the curated published datasets were converted to gene expression TPM values and integrated with our gene expression data. To remove technical batch effects, we applied the *removeBatchEffect* function from the *limma* package (version 3.58.1). We then compared the first two principal components of the integrated datasets to assess whether they exhibit similar transcriptome profiles.

### Differential expression analysis

Counts from expressed transcripts across all cell types were used to perform differential gene expression (DGE) and differential transcript expression (DTE) analysis using the DESeq2 (version-1.42.1) R package. Pairwise differential tests were conducted between cell types, such as fibroblasts vs. iPSC, iPSC vs. iCN, and fibroblasts vs. iCN. Genes and transcripts were considered to have differential expressions between cell types if their corresponding test adjusted P-values were ≤ 0.05 and |log2FC| > 1.

Upregulated genes and transcripts with significant DGE or DTE events (log2FC > 1) were selected for Gene Ontology (GO) term enrichment tests. GO term tests were performed using gProfiler2(v0.2.3) in R and the top 10 GO terms with the best adjusted p-values from each category—biological processes (BP), cellular components (CC), and molecular functions (MF) were selected for visualization.

### Differential transcript usage analysis

Differential transcript usage (DTU) analysis was performed using the IsoformSwitchAnalyzeR v2.2.0 framework in combination with DRIMSeq (Vitting-Seerup & Sandelin, 2019; Nowicka & Robinson, 2016). DRIMSeq conducted the DTU analysis based on transcript usage calculated from TPM values of expressed transcripts.

We integrated the DRIMSeq results with gene and transcript differential expression results produced by DESeq2 (in Differential expression analysis section) within IsoformSwitchAnalyzeR framework to predict alternative splicing events and their consequences. Additionally, to ensure robust results, we applied several filtering steps as recommended by IsoformSwitchAnalyzeR. Initially, transcripts were filtered based on a minimum gene expression threshold (gene_condition_1 > 1 and gene_condition_2 > 1) to exclude noise. DTU events were identified using DRIMSeq test results with an adjusted p-value < 0.05 and an absolute usage difference (dIF) > 0.1. Identified DTU genes were then selected for Gene Ontology (GO) term enrichment tests. gProfiler2(v0.2.3) in R and the top 10 GO terms with the best adjusted p-values from each category—biological processes (BP), cellular components (CC), and molecular functions (MF) were selected for visualization.

To predict the functional consequences of differentially utilized transcripts, we analyzed coding potential (CPC2) (Y. J. Kang et al., 2017), protein domains (Pfam) (Mistry et al., 2021), signal peptides (SignalP 5.0) (Nielsen et al., 2024), intrinsically disordered regions (IDR) (Babu, 2016), using external sequence analysis tools including Pfam, CPC2, SignalP-5.0, and IUPred2A (Mészáros et al., 2018).

## List of Abbreviations

AD: Alzheimer’s disease
APP: Amyloid precursor protein
AS: Alternative splicing
BP: Biological processes
BSCL2: Berardinelli-Seip Congenital Lipodystrophy 2
CC: Cellular components
CDS: Coding sequence
DAI: Days after induction
DGE: Differential gene expression
DTE: Differential transcript expression
DTU: Differential transcript usage
FN1: Fibronectin-1
HD: Huntington’s disease
HSP: Hereditary Spastic Paraplegia
iCN: iPSC-derived cortical neuron
iPSC: induced pluripotent stem cell
KIF2A: Kinesin Superfamily Protein 2A
KPI: Kunitz-like protease inhibitor
LR: Long-read
lncRNA: long non-coding RNA
MF: Molecular functions
OMIM: Online mendelian inheritance in man
PD: Parkinson’s disease
PCA: Principal component analysis
PSEN1: Presenilin 1
RBP: RNA-binding protein
RNA-seq: RNA-sequencing
TPM: Transcripts per million

## Author’s Contributions

R.S., J.X., M.H. and M.N. designed the study; M.H., K.M., M.N., and S.H. established the cell lines and prepared the cells for sequencing; E.B-A., J.A., S.O. and N.C. performed long-read library, sequencing and primary analysis for amplified cDNA. ; R.S., J.X., and M.H. analyzed the data; E.B.A., R.S., J.X. and M.H. wrote the paper with input from all authors. All authors critically revised and acknowledged the final version of the manuscript.

## Acknowledgment

We are grateful to Yvonne Schelling, Stefanie Schuster, Philip Höflinger and Milena Korneck for their assistance in generating, characterizing, culturing and differentiating some of the cell lines used in this study and to Ludger Schöls for his contribution to obtaining some of the proband samples. We thank Noella-Theresa Ansorge for the assistance in library preparation and sequencing.

This study was funded by the Horizon 2020 research and innovation program (grant 779257 ‘Solve-RD’ to RS), the Bundesministerium für Bildung und Forschung (BMBF) through funding for the TreatHSP network (grant 01GM2209A to RS), the National Institute of Neurological Disorders and Stroke (NINDS) and the National Institutes of Health (NIH) under Award Number R01NS072248 (RS), and the Clinician Scientist Programme PRECISE.net funded by the Else Kröner-Fresenius-Stiftung (RS). RS is a member of the European Reference Network for Rare Neurological Diseases – Project ID 739510. NGS sequencing methods were performed with the support of the DFG-funded NGS Competence Center Tübingen (INST 37/1049-1).

